# PSEN1^M146V^ and PSEN1^A246E^ mutations associated with Alzheimer’s disease impair proper microglia differentiation

**DOI:** 10.1101/2023.10.08.561397

**Authors:** Antoine Aubert, Maria Grazia Mendoza-Ferri, Aude Bramoulle, François Stüder, Bruno Maria Colombo, Marco Antonio Mendoza-Parra

**Affiliations:** UMR 8030 Génomique Métabolique, Genoscope, Institut François Jacob, CEA, CNRS, University of Evry-val-d’Essonne, University Paris-Saclay, 91057 Évry, France

**Author notes:** Corresponding author: Marco Antonio Mendoza-Parra. These authors contributed equally (Bruno Maria Colombo, Marco Antonio Mendoza-Parra).

**Keywords:** Alzheimer’s disease, PSEN1, microglia

## Abstract

Genetic variants associated with the late onset of Alzheimer’s disease (AD), were correlated with genes known to be expressed in microglia, suggesting for an AD-genetic component directly influencing microglia behavior. Instead, the role of the familial AD (fAD) genetic mutations was systematically studied from the angle of the Amyloid-Beta pathway; leaving their participation in microglia homeostasis unexplored.

Here we demonstrate that two previously described fAD-related PSEN1 mutations directly impair proper microglia differentiation. While human induced pluripotent stem cells harboring the PSEN1-M146V mutation did not give rise to hematopoietic precursor (HPC) intermediate during microglia differentiation, a PSEN1-A246E mutant line managed to produce HPCs, but died within the first days of microglia differentiation.

Detailed transcriptomics/epigenomics and functional assays revealed the setup of a pro-apoptotic program in the PSEN1-A246E mutant, which was circumvented when HPCs were grafted in brain organoids (BORGs). Microglia obtained in BORGs presented preferentially pro-inflammatory signatures, further supported by their correlation with recent data providing a detailed stratification of the various microglia populations within AD-patient samples.

Overall, this study contributes to reconsider the influence of the previously identified familial mutations in the homeostasis of this immune component of the central nervous system.

---

Alzheimer’s disease (AD) is a progressive neurodegenerative disorder that impairs memory, behavior and thinking ^1^ . From a molecular angle, AD is characterized by the extracellular accumulation of beta amyloid plaques (Aβ) and by the presence of neurofibrillary tangles due to the intraneuronal aggregation of hyperphosphorylated Tau protein ^2^. These two major hallmarks were considered over years as the source of the progressive neurodegeneration, notably due to the fact that microglia – the brain’s immune cells – accumulate around the Aβ plaques, as an inflammatory response ^3–5^. Nevertheless, it is known that microglia are also involved in the generation of a vicious inflammatory cycle inducing Aβ plaques accumulation and the release of inflammatory mediators ^6–8^, phenomena associated with the worsening of the disease.

Genome-wide association studies (GWAS) – revealing single-nucleotide polymorphisms (SNPs) variants associated to late onset of the disease (LOAD) – were recently correlated with genes known to be specifically expressed in microglia, strongly arguing for an AD-genetic component directly influencing microglia behavior ^9–11^. In contrary, the influence of the familial AD (fAD) genetic mutations (found in the amyloid precursor protein (APP) and presenilin (PSEN1 and PSEN2) genes) in microglia homeostasis remained unexplored till now.

To explore this potential genetic influence, we have first implemented the method described by A. McQuade et al ^12^, to generate human microglia from induced pluripotent stem cells (hiPSCs), by passing through the intermediate of hematopoietic precursor cells (HPCs) (**Fig.1A**). This protocol allowed us to obtain HPC and microglia cells (iMGL) with yields > 95%, as revealed by flow cytometry (**Fig. 1B&D & Extended Data Fig.1**), further validated by quantitative RT-PCR targeting specific markers for both HPCs (CD34, CD43, CD45; **Fig. 1C**) and iMGL (CD11b, TREM2, TMEM119, IBA1; **Fig. 1E**). In addition to this molecular characterization, we have validated the functional response of the obtained microglia, by their uptake capacity of fluorescent latex beads (**Fig. 1F**). To fully characterize microglia differentiation, we have generated bulk transcriptomes for HPC intermediate but also for three different time-points during microglia formation (19, 26 and 41 days). Comparing their gene-expression signatures with those assessed from publicly available datasets (Principal component analysis (PCA)), revealed strong similarities with both *in vitro*-generated microglia, as well as those collected from brain samples (**Fig. 1G**).

**Fig 1.**
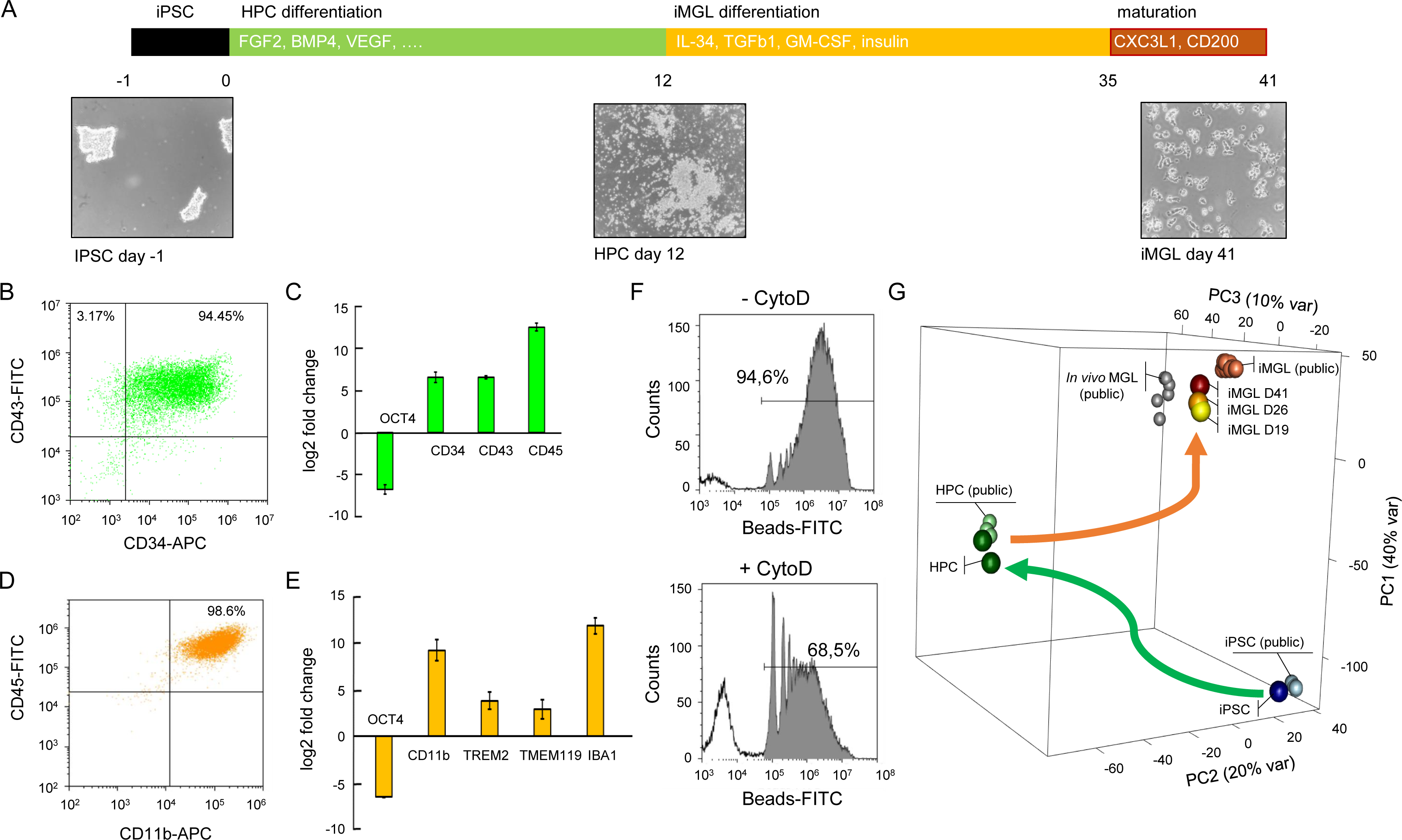
Differentiation of human induced pluripotent stem cells (iPSC) into microglia. **A.** Schematic representation of the protocol used for generating induced microglia-like (iMGLs) cells by the intermediate of hematopoietic progenitors (HPC). Human iPS cell colonies seeded as small aggregates (day −1) generated large numbers of round HPCs (day 12). Harvested HPCs were first treated with three cytokines (IL-34, TGFB, GM-CSF, then exposed to two maturation factors (CXC3L1, CD200); leading to iMGLs (day 41). **B.** Flow cytometry analysis of HPCs (day 12), based on CD34 and CD43 expression, revealing a pure population at ∼95%. **C.** Quantitative RT-PCR assay in HPCs, targeting genes coding for the hematopoietic markers CD34, CD43, CD45 and the pluripotent marker OCT4. **D.** Flow cytometry analysis of iMGL (day 41), based on CD11b and CD45 expression. **E.** Quantitative RT-PCR assay in iMGL, targeting the microglial gene markers CD11b, TREM2, TMEM119, IBA1 and the pluripotent marker OCT4. **F.** Flow cytometry assay to assess phagocytosis of fluorescent latex beads by iMGLs in presence or absence of Cytochalasin D, a potent inhibitor of actin polymerization, as a control for phagocytosis. **G.** 3D principal component analysis displaying global transcriptome assays performed in iPSCs, HPCs and three stages during iMGL differentiation (day 19, day 26, day 41), in comparison with public datasets ^12^.

This validated protocol has been applied to hiPSCs harboring either the familial APP Swedish (Swe), the PSEN1-M146V or the double mutant APP Sew/PSEN1-M146V relative to their isogenic wild-type (WT) control line. Surprisingly, while both WT and the APP Swe mutant line reached the HPC intermediate stage, both PSEN1-M146V and PSEN1-M146V/APP Swe isogenic mutant lines failed to reach this stage (**Fig. 2A & Extended Data Fig.2A**). Further differentiation of both WT and the APP Swe mutant line gave rise to fully differentiated microglia, confirmed by the presence of CD11b CD45 double positive cells (**Fig. 2B & Extended Data Fig.1 & 2A**), gene expression signatures assessed by RT-qPCR, and bulk transcriptomics (**Extended Data Fig.2**). Obtained microglia from both WT and the corresponding isogenic APP Swe mutant line presented similar uptake phagocytosis-like behavior and inflammatory response signatures in presence to LPS (**Fig. 2C&D**).

**Fig 2.**
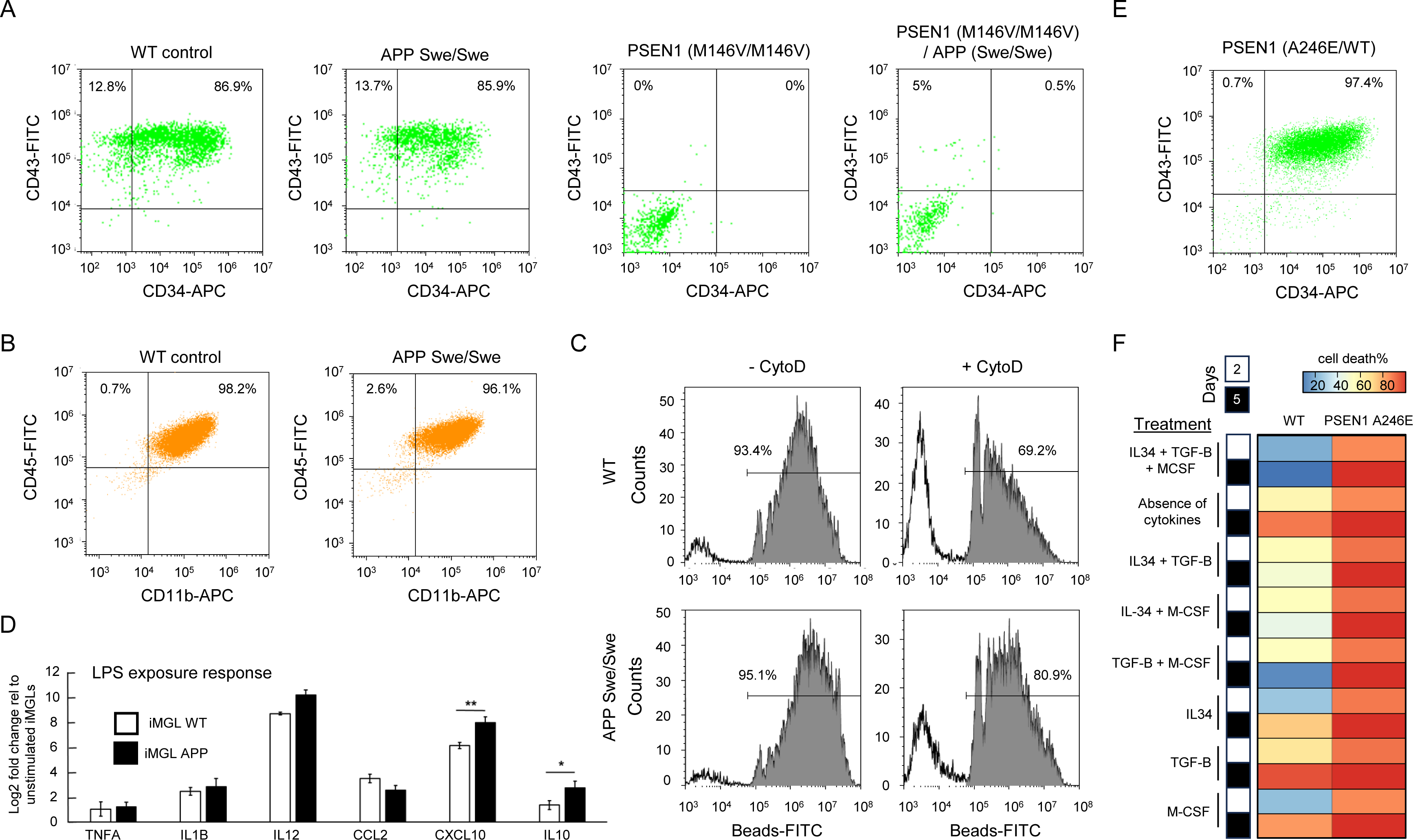
Microglia differentiation from human IPSCs harboring autosomal-dominant AD-related mutations. **A.** Flow cytometry analysis of different isogenic HPCs (WT, APP Swe/Swe, PSEN1 M146V/M146V and APP Swe/Swe/PSEN1 M146V/M146V), based on CD34 and CD43 expression. Notice that HPC differentiation was only possible for the WT control and the APP Swe/Swe mutant line. **B.** Flow cytometry analysis of isogenic iMGLs (WT and APP-Swe/Swe, respectively), based on CD11b and CD45 expression, confirming a pure microglial population at >96%. **C.** Flow cytometry revealing the uptake of fluorescent latex beads by isogenic iMGLs (WT and APP-Swe/Swe, respectively) in presence or absence of Cytochalasin D as control of phagocytosis. **D.** Liposacharide (LPS)-driven inflammation response assessed in isogenic iMGLs (WT and APP-Swe/Swe). Gene expression induction associated to cytokines and chemokines secretion after LPS stimulation was assessed by quantitative RT-PCR assay. **E.** Flow cytometry analysis performed in HPCs harboring the PSEN1-A246E mutation (day 12), confirming a pure population at >97%. **F.** Apoptosis-mediated cell death of HPCs harboring the PSEN1-A246E mutation during the iMGL differentiation. HPCs were treated with the three-cytokines cocktail, required for initiating microglia differentiation (IL-34, TGF-b, M-CSF), or in presence or absence of one or two of them, and assessed by Flow cytometry (2 days and 5 days after treatment) targeting Anexin V and Propidium Iodide.

The astonishing observation that, in contrast to their isogenic counterparts, the PSEN1-M146V mutant lines did not reach even the HPC intermediate state, argued for a major role of this gene in microglia ontogenesis. To explore this hypothesis, we took advantage of another human iPSC line, issued from an AD patient, and harboring a different PSEN1 mutation, namely PSEN1-A246E. Interestingly, PSEN1-A246E mutant hIPSCs gave rise to >97% CD34 CD43 positive cells, characteristic of HPCs (**Fig. 2E**). Confident with this success, we have attempted to generate iMGLs from this HPC intermediate, but cells died during the first 5 days of differentiation (**Extended Data Fig.3A**). Since microglia differentiation is driven by the exposure to three cytokines (IL-34, TGF-Beta and M-CSF), we tried to elucidate the potential deleterious effect of these factors during the differentiation of PSEN1-A246E. Indeed, Annexin V and Propidium Iodide (PI) staining revealed >80% of apoptotic cells from the second day of treatment with any combination of the aforementioned cytokines (**Fig. 2F & Extended Data Fig.3B**).

Puzzled by the fact that HPCs harboring the PSEN1-A246E mutation perishes during the first days of microglia differentiation, we tried to reveal the difference of these cells with those issued from WT control lines. Bulk transcriptomes revealed similar number of differentially regulated genes in both HPCs relative to their corresponding hiPSCs (∼880 up-regulated genes and ∼1,200 down-regulated genes; **Fig. 3A**). Furthermore, 592 genes were found commonly up-regulated, while 282 genes were specifically up-regulated in WT HPCs, and other 293 genes were induced specifically in HPCs harboring the PSEN1-A246E mutation (**Extended data Fig.4 & Fig. 3B**). Interestingly, the specifically up-regulated genes found in WT HPCs were preferentially enriched for the survival pathways WNT, VGEF (signaling factor in mediating chemotactic responses of immune responding cells ^13^) and PI3K-Akt signaling (involved in the expression and production of pro-inflammatory mediators^14^), while those associated to HPCs harboring the PSEN1-A246E mutation were enriched for TNF signaling pathway, suggesting a pro-inflammatory and proapoptotic state (**Fig. 3B**).

**Figure 3.**
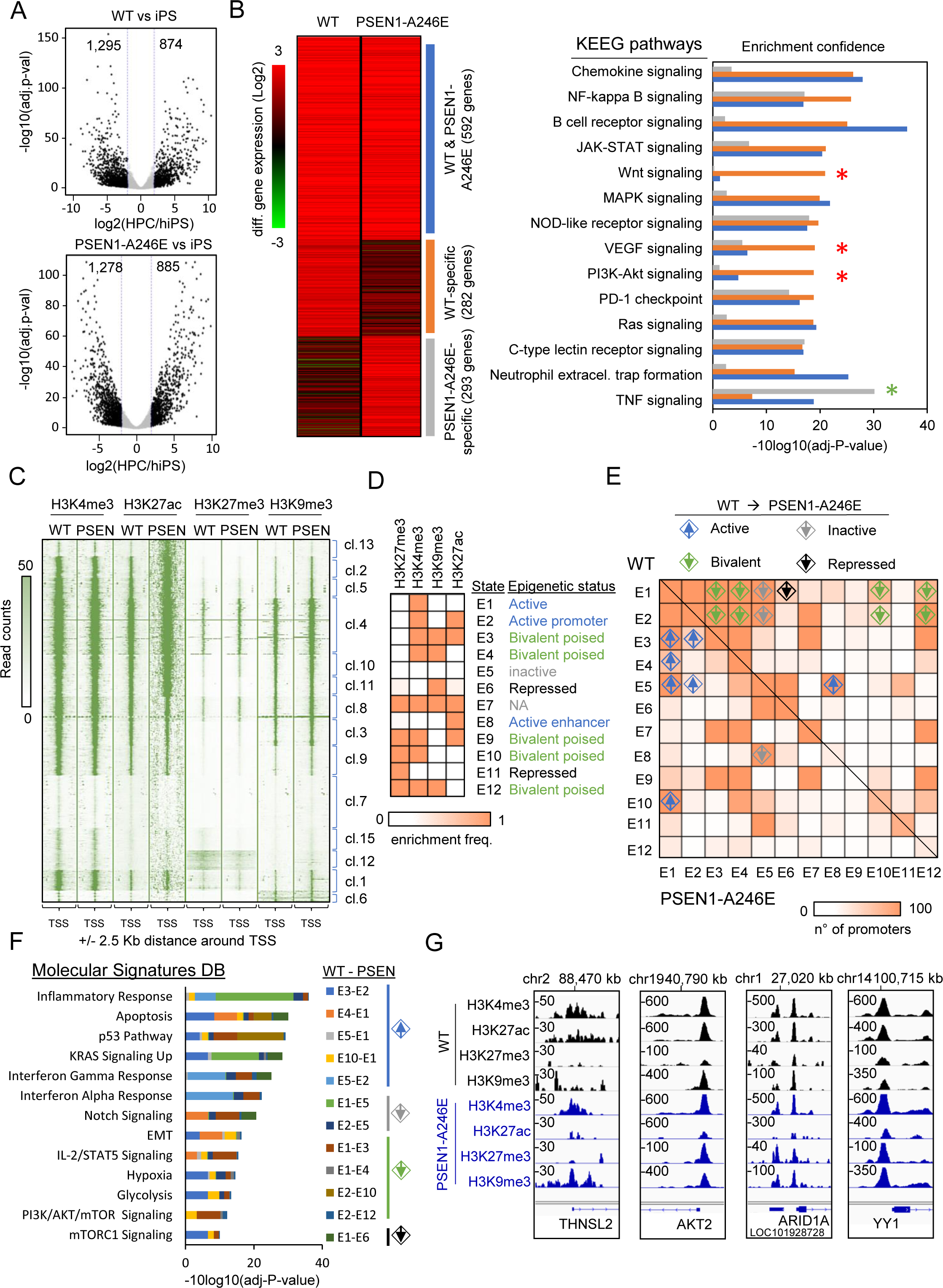
Transcriptomics and epigenomics profiling performed in WT and PSEN1-A246E mutant HPCs reveal a deregulation of the inflammatory and apoptotic programs. **A.** Volcano plot displaying the number of differentially expressed genes in either WT (top panel) or PSEN1-A246E mutant HPCs relative to their corresponding iPSCs. **B.** Heatmap displaying the number of common or specific upregulated genes between the WT and the PSEN1-A246E mutant HPC line (log2 foldchange>3; left panel), and their corresponding enrichment within KEEG pathways (right panel). Notice the specific enrichment of the Wnt, VEGF and PI3K-Akt signaling pathway for genes specifically upregulated in WT HPCs (red star), and the TNF signaling pathway for those specific in PSEN1-A246E mutant HPCs (green star). **C** Histone modifications enrichment around gene promoters (+/-2.5 Kb) assessed in both HPC lines. 15 different classes were identified (hierarchical clustering; cl) on the basis of the read count enrichment levels across different histone modifications. **D.** Combinatorial chromatin state analysis revealing functionally relevant promoter status within HPC cells. **E.** Chromatin state transitions between WT and PSEN1-A246E mutant HPCs highlighting promoter chromatin status changes (Active, Inactive, bivalent poised and repressed). **F.** Relevant pathways affected by the promoter chromatin status changes (Molecular signature database). **G.** Example of promoter chromatin status changes between WT and PSEN1-A246E mutant HPCs.

**Fig 4.**
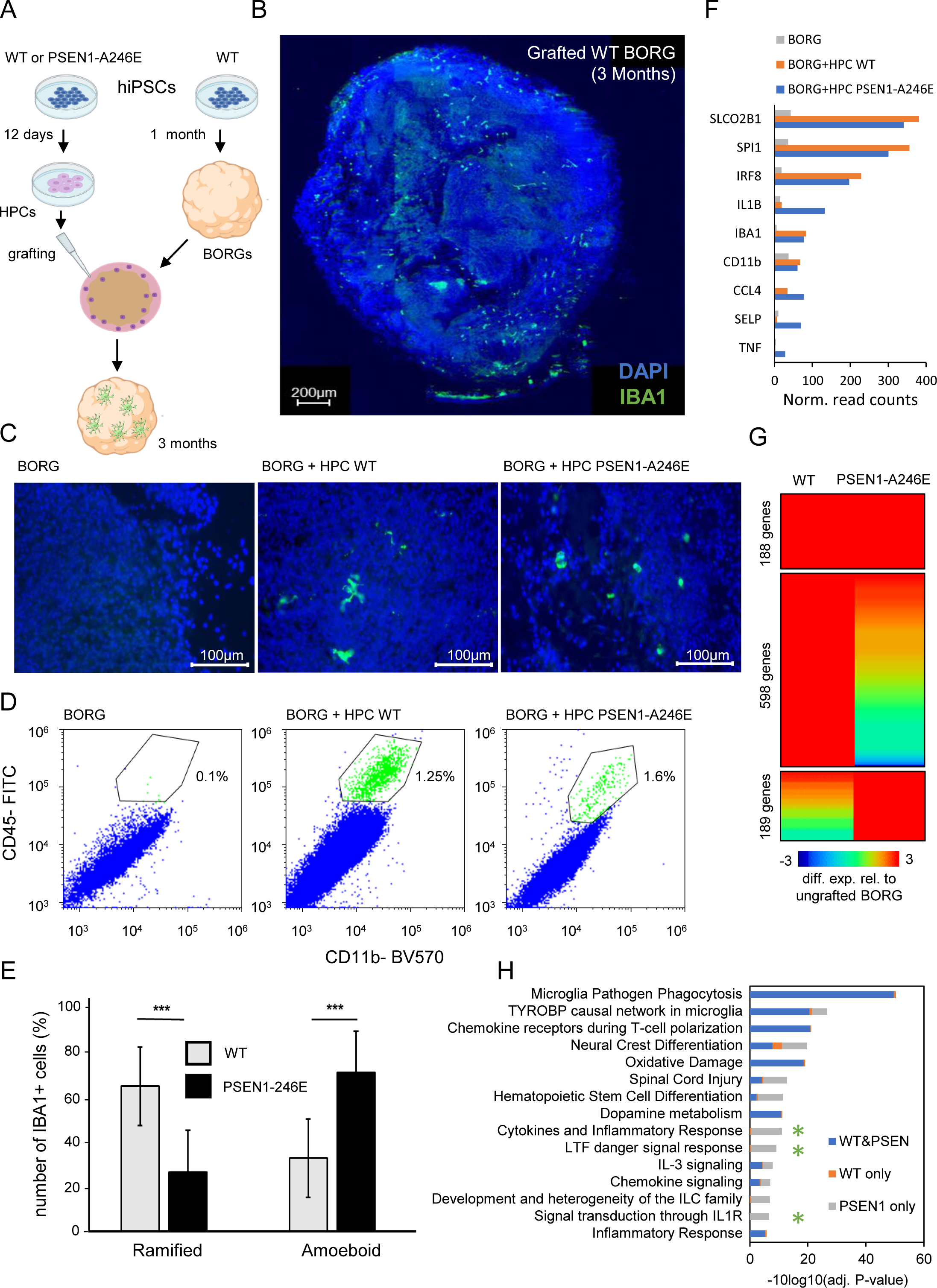
HPCs harboring the PSEN1-A246E mutation differentiated into microglia when grafted into human brain organoids (BORGs) **A.** Scheme illustrating the strategy in use for grafting HPCs into 1-month old BORGs. 12-days differentiated HPCs are embedded in Matrigel together with a WT BORG, then cultured during 2 months more leading to microglia differentiation. Scheme generated with <u>Biorender</u>. **B.** Immunofluorescence micrograph of a WT BORG grafted with WT HPCs, labelled for the microglia marker IBA1 (green). Nuclei were revealed by DAPI (blue). Micrograph has been obtained by manual scanning across the BORG section. **C.** Immunofluorescence micrograph of ungrafted (left) and grafted BORGs with either WT (middle) or PSEN1-A246E mutant HPCs (right), labelled for the microglia marker IBA1 (green). Nuclei were revealed by DAPI (blue). Notice the differences in shape between the IBA1+ cells observed in the WT (ramified-like) and PSEN1-A246E grafted HPCs (round amoeboid). **D.** Flow cytometry assay targeting the CD11b and CD45 markers in the aforementioned ungrafted or grafted conditions. The indicated double-positive labelling revealed >1% of microglial cells within BORGs issued from grafted HPCs. **E.** Fraction of IBA1+ cells (in percent), assessed in grafted BORGs with either WT or PSEN1-A246E HPCs, presenting either ramified or round-amoeboid-like shapes, indicative for resting or activated microglial state. **F**. Normalized read-counts observed in microglial gene markers obtained from bulk transcriptomics performed in ungrafted (BORG), or grafted BORGs with the aforementioned HPCs. **G.** Differential gene expression analysis (log2 fold-change>3) performed between grafted samples and the ungrafted BORG control, revealing the number of up-regulated genes in common or specific for each grafted condition. **H.** Cumulative Pathway enrichment analysis performed for each of the up-regulated group of genes described in G. Green stars highlight relevant enriched pathways associated to upregulated genes specific to BORGs grafted with PSEN1-A246E HPCs.

To further understand the differences between HPCs issued from WT or PSEN1-A246E mutant iPSC lines, we have explored their chromatin epigenetics status by targeting the histone modifications H3K4me3 (commonly retrieved in transcription start sites; TSS), H3K27ac (retrieved on both active enhancers and promoters) and the repressive marks H3K27me3 and H3K9me3 (**Fig. 3C**). This analysis revealed promoter regions strongly enriched for repressive marks in both HPC lines (cluster 6: H3K9me3; cluster 12: H3K27me3; **Fig.3C**), but also others presenting either active promoter markers (cluster 13), or bivalent chromatin promoter status, including the presence of H3K9me3 together with the H3K4me3 (e.g. cluster 4,10,11,1; **Fig. 3C**) as described in previous studies ^15^. A combinatorial epigenetic status analysis allowed to better identify the presence of these different chromatin states within both lines (**Fig.3D**), but also identify promoter status differences between WT and PSEN1-A246E mutant HPCs. Promoters were broadly classified in four groups: (i) those that gain active signatures in PSEN1-A246E line relative to WT; (ii) those that got inactivated; (iii) repressed; or (iv) those that reached a bivalent poised state (**Fig. 3D&E**). Importantly, enrichment pathway analysis performed over most of the relevant promoter transitions revealed the inactivation of the inflammatory response pathway, a gain in apoptotic signatures as well as in the interferon Gamma & interferon Alpha response (**Fig. 3F**), suggesting for the setup of a counteracting epigenetics activity against the observed pro-inflammatory status of the system.

Indeed, promoters associated to genes like THNSL2 (inducing the inflammatory cytokine IL-6 production ^16^), AKT2 (coding for a member of the family of AKT kinases, known to regulate processes like cell proliferation and survival), ARID1A (coding for a component of the neural progenitors-specific chromatin remodeling complex (npBAF complex) and the neuron-specific chromatin remodeling complex (nBAF complex), essential for the self-renewal/proliferative capacity of the multipotent neural stem cells), or YY1 (coding for the transcriptional repressor protein YY1, recently shown to be an essential regulator of hematopoietic stem cell quiescence ^17^) are retrieved in a bivalent poised state in HPCs harboring the PSEN1-A246E mutation; suggesting a deregulation of survival pathways (**Fig. 3G**). Altogether, these results suggest that WT HPCs have a phenotype prone to survival which would allow them to easily differentiate into microglia. On the contrary, HPCs harboring the PSEN1-A246E mutation present a deregulated epigenome encoded pro-inflammatory / apoptotic response program, leading to death within the first days of exposure to the cytokinic environment required for microglia differentiation.

The fact that HPCs harboring the PSEN1-A246E mutation are not able to reach microglia differentiation is contradictory with the observation of microglia in AD-patients harboring this and other PSEN1 mutations ^18–20^. For this reason, we implemented a microglia differentiation strategy closer to the in vivo situation; i.e. within a complex system harboring a variety of neurons and glial cells. Specifically, we have grafted HPCs in young human brain organoids (1-month-old BORGs) with the expectation that this complex 3-dimensional nervous tissue will promote microglia differentiation (**Fig. 4A**). BORGs grafted with WT HPCs gave rise to IBA1-positive cells after two months, and the same has been observed when BORGs were grafted with HPCs issued from PSEN1-AD246E mutant line (**Fig. 4B&C and Extended data Fig.5**). Flow cytometry revealed comparable numbers of CD11b CD45 double positive cells in grafted BORGs with HPCs issued from WT or PSEN1 mutant line (∼2% of cells), suggesting for a similar performance of microglia differentiation within BORGs (**Fig. 4D**). Interestingly, IBA1+ cells issued from the PSEN1 mutant line presented up to 70% of round-body shaped cells rather than the ramified structures, preferentially observed in the WT counterpart; indicative for the presence of pro-inflammatory amoeboid microglia issued from the PSEN1 mutant line (**Fig. 4C&E**).

**Figure 5.**
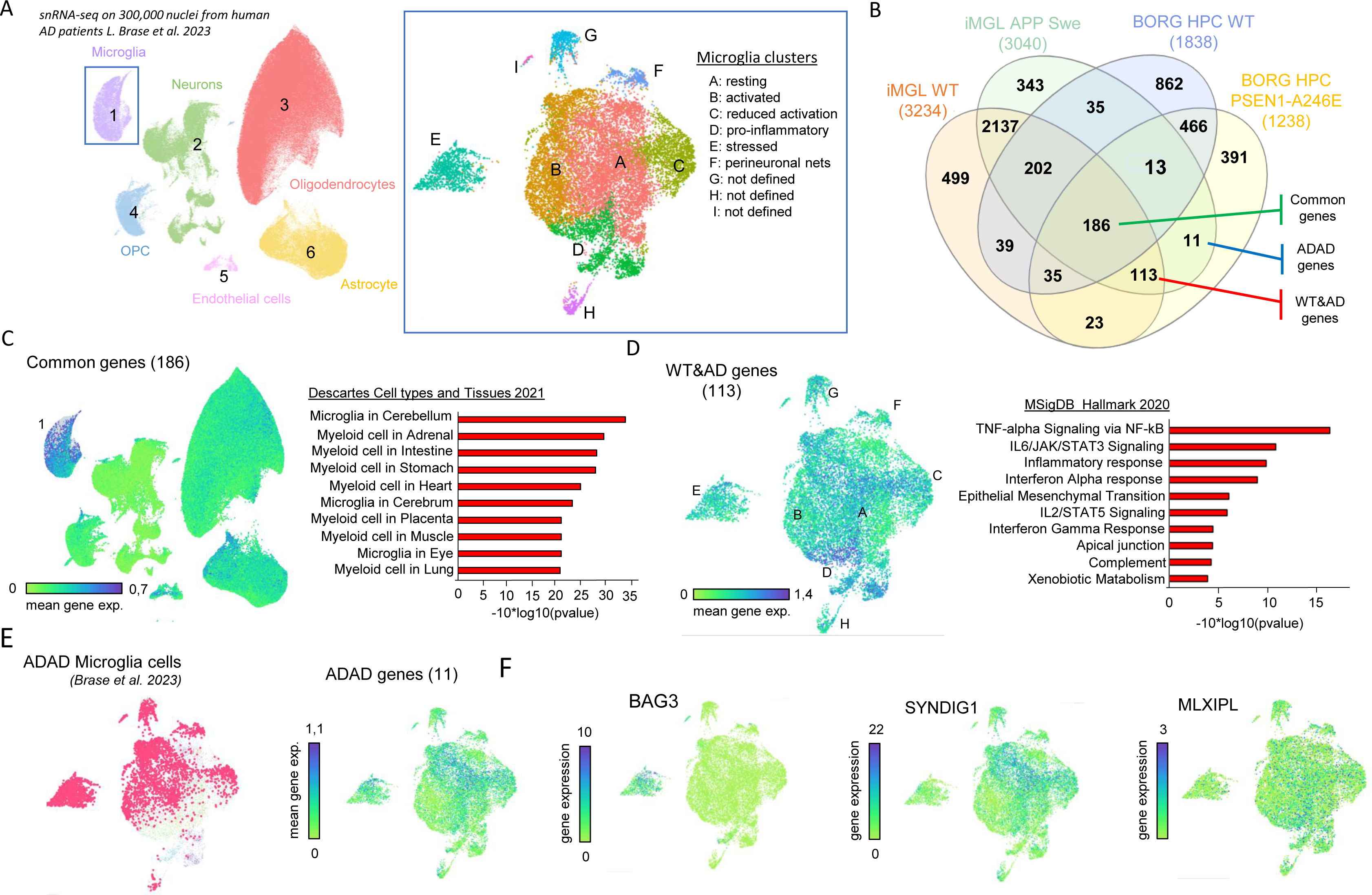
Microglial signatures assessed by HPC differentiation in-vitro present a strong correlation with those retrieved in a single-nucleus transcriptome atlas issued from AD patients. **A.** UMAP plot displaying six major clusters assessed from the ∼300,000 single-nuclei RNA-sequencing atlas generated from the parietal cortex of AD autosomal dominant (APP and PSEN1) and risk-modifying variant (APOE, TREM2, MS4A) carriers ^18^ . The Inset (right panel) displays a detailed stratification of the microglial cluster (1), as described by Brase and colleagues. **B.** Venn-diagram illustrating the number of up-regulated genes in four microglial differentiation conditions described in this study. Three major groups of interest are highlighted: Upregulated genes common to all four cases (“common” genes: 186); Upregulated genes specific to the WT and APP Swe/Swe mutant microglia as well as to those obtained by grafting PSEN1-A246E mutant HPCs in BORGs (herein defined as “WT&AD” genes: 113); those specific to the APP Swe/Swe and the PSEN1-A246E mutant background (herein named as “ADAD” genes, for Autosomal Dominant AD: 11). **C.** Left panel: Mean gene expression signature associated to the in-vitro up-regulated “common” genes across all single-nuclei UMAP plot, revealing a strong enrichment in the microglial cluster (1). Right panel: Cell types and Tissues signature enrichment analysis performed in the “common” genes, confirming the microglial signature. **D.** Left panel: Mean gene expression signature associated to the in-vitro up-regulated “WT&AD” genes across all single-nuclei UMAP plot, revealing a strong enrichment in the microglial resting (A), pro-inflammatory (D) and reduced activation (C) clusters. Right panel: Pathway enrichment analysis performed in the “WT&AD” genes, confirming the presence of inflammatory response signatures. **E.** Left panel: Autosomal Dominant AD (ADAD)-associated cells within the microglial cluster issued from the single-nuclei RNA-sequencing atlas. Right panel: Mean gene expression signature associated to the in-vitro up-regulated “ADAD” genes across the same microglial cluster. **F.** Gene expression signature within the microglial cluster assessed for the genes BAG3, SYNDIG1, and MLXIPL, retrieved within the 11 upregulated “ADAD” genes, previously described to be upregulated in microglia in AD context.

Microglia formation in grafted BORGs has been further confirmed by RT-qPCR (**extended data Fig.5D**) and bulk transcriptomics, revealing the presence of gene markers like SLCO2B1 (coding for a microglia-enriched plasma membrane heme importer ^21^), SPI1 (coding for the transcription factor PU.1, specifically expressed in microglia in the brain, and considered as a major player of the core transcription regulatory circuitry defining microglia cell identity ^22^), IRF8 (Interferon regulatory factor 8, known to be essential for proper microglia development and reactivity ^23,24^) or IBA1/AIF1 (coding for a calcium-binding protein, known to participate in intracellular processes like phagocytosis, membrane ruffling and F-actin remodeling) (**Fig. 4F**). A differential gene expression analysis relative to ungrafted control BORGs revealed 188 genes induced in both BORGs grafted with either WT or PSEN1-A246E HPCs, 598 up-regulated genes specific to BORGs grafted with WT HPCs, and 189 genes induced specifically in BORGs grafted with PSEN1-A246E HPCs (**Fig. 4G**). Importantly, common upregulated genes were enriched in pathways associated to Microglia Pathogen Phagocytosis, or TYROBP causal network microglia pathway (microglial-enriched phosphotyrosine protein, genetically associated with late onset Alzheimer’s disease ^25^). Interestingly, genes specifically upregulated in BORGs grafted with PSEN1-A246E HPCs were enriched for cytokines and inflammatory response pathways (LTF danger signal response pathway, Signal transduction though IL 1R), strongly supporting for their over-reactive proinflammatory status (**Fig. 4H**).

Finally, to evaluate the relevance of microglia signatures retrieved in our in-vitro study with readouts assessed form AD-patient related samples, we have compared these results with a recent study covering ∼300 thousand nuclei from the parietal cortex of either inherited genetic AD-patients (presenting mutations in *APP* and *PSEN1*) or presenting genetic risk factor variants (APOE, TREM2 and MSE4A) ^18^ (**Fig.5A**). For this, we have first compared the different type of microglia like cells (iMGL) generated in this study on the basis of their up-regulated genes; namely: microglia issued from WT hiPSCs (iMGL WT), those issued from the APP Swe/Swe mutant iPSCs (iMGL APP Swe), microglia obtained by grafting WT HPCs as well as those obtained by grafting HPCs harboring the PSEN1-A246E mutation (**Fig.5B**). Among all shared up-regulated genes (**Extended data Fig.6**), we have focused our attention to three major groups: 186 commonly upregulated genes by all aforementioned conditions (herein described as “common” genes); 113 genes shared within the iMGL WT, the iMGL APP and the BORGs HPC PSEN1-A246E (herein described as “WT&AD” genes); and 11 genes specifically over-expressed by iMGL cells issued from hiPS cells harboring AD-related mutations (herein defined as “ADAD” genes” for autosomal dominant AD mutations as described by Brase et al ^18^) (**Fig. 5B**). Each of these shared up-regulated genes were compared with the single-cell transcriptome signatures stratified in the study of Brase et al ^18^, revealing that common upregulated genes within all in-vitro generated microglia are over-represented in the microglia cluster issued from AD-patients samples (**Fig.5C**). Similarly, upregulated genes denoted as “WT&AD” are strongly enriched within the microglia cluster issued from AD-patients, and notably with microglia cells labelled as pro-inflammatory (cluster D), in agreement with the presence of TNF-alpha signaling pathway enrichment (**Fig. 5D**). Finally, the ADAD group of genes presented a strong correlation with the “ADAD” microglia cells characterized by Brase et al (**Fig.5E**), and notably supported by the presence of the genes BAG3 (chaperone protein, described as essential for Tau and APP degradation via phagocytosis ^26^), SYNDIG1 (considered as a central regulator of excitatory synapse development ^27^, but also described as being over-expressed in microglia ^28^), or MLXIPL (a glucose-responsive transcription factor also known as ChREBP), a gene for which previous GWAS studies revealed significant association of single nucleotide polymorphisms (SNPs) in the context of AD ^29^. SYNDIG1 and MLXIPL were also identified in a recent single-nuclei RNA-seq study covering ∼120,000 microglia isolated from 12 AD and 10 control donors ^30^ (**Fig.5F**).

While previous studies tempted to explain the autosomal dominant AD mutations by their direct role in the beta-amyloid processing pathway taking place within neurons, herein we demonstrate that the aforementioned PSEN1 mutations directly impair proper microglia differentiation. Two PSEN1-M146V mutant hiPS lines did not even manage to reach the HPC state, while the PSEN1-A246E mutant line died within the first days of microglia differentiation. Our in-depth epigenetics and transcriptomics analysis, revealed the setup of a pro-apoptotic program in the PSEN1-A246E mutant line, which appeared to be circumvented when HPCs were grafted in brain organoids. Nevertheless, the obtained microglia presented preferentially pro-inflammatory signatures, which fully correlated with a recent study providing a detailed stratification of the various microglia populations within AD-patient samples ^18^.

In essence, this study contributes to reconsider the role of microglia in AD, and notably to reevaluate the influence of the previously identified autosomal dominant mutations in the homeostasis of this immune component of the nervous system.

## Materials & Methods

### iPSCs origin, amplification, and maintenance

Human induced pluripotent stem cells (iPSCs) used in this study were obtained from the New York stem cells foundation (Wild-type parental iPS line 7889SA: CO0002-01-SV-003; isogenic Knock-in constructs: APPswe / APPswe: CO0002-01-CS-004 / BN0013; PSEN1-M146V/PSEN1-M146V: CO0002-01-CS-001; double Knock-in APPswe/APPswe & PSEN1-M146V / PSEN1-M146V: BN0002) and from the centre of Regenerative Medecine in Barcelona – Banc de Linies Cellulars (CRMB) (Wild-type iPS line Ctrl2-R4F-5; PSEN1-A246E / WT). IPSCs were amplified in CytoOne 6 well-plate culture plates (Starlabs, 2550254) coated with hESC qualified matrigel (Corning, 354277), in mTeSRplus (StemCell Technologies, 100-0276) at 37°C; 5%CO_2_. IPSC medium was changed every two days and was passaged once a week when confluency reached around 80% using ReleasR (StemCell technologies, 05872, 100-0484) solution.

### Differentiation of microglia-like (MGL) cells from iPSCs

Microglia differentiation has been performed in two steps, following the protocol described by McQuade et al 2018 ^12^. Briefly, iPSCs were first cultured with STEMDiff Hematopoietic kit (StemCell technologies, 05310) for 12 days in a CytoOne 12-well plate (Starlabs, 2550253) coated with hESC qualified Matrigel to generate HPCs. Floating CD43+ positive cells were transferred in a 6-well plate (100,000 cells/well) coated with hESC qualified matrigel and cultured in microglial differentiation medium (DMEM/F12 Life technology, 11330057), 1X MEM-NEAA (Life technology, 11140050), 1X GLUTAMAX (Life technology, 35050-038), 2X Insulin-Selenium transferrin solution (Life technology, 41400045), 5µg/mL Insulin (Sigma-Merck, I2643-50MG),0.5X N2 (Life technology, 17502048), 2X B27 (Life technology, 17502048), and 400µM Monothioglycerol (Sigma-Merck, M1753-100ML)). Microglia differentiation was promoted by supplementing medium with three cytokines:IL-34 (100ng/mL; Peprotech, 200-34), MCSF (25ng/mL; Milteny, 130-096-489), and TGF-B1 (50 ng/mL; Milteny, 130-095-066). 2mL of this supplemented medium were added at the first day, then 1mL were added every two-days. When the volume reached 7mL, 6mL containing cells was centrifuged (5 minutes, 1100 rpm), pelleted-cells were resuspended in 1 mL of fresh medium and replaced in the plate containing the remaining 1 mL. In addition to the aforementioned cytokines, two new others were added after day 26 of microglia differentiation: CD200 (100 ng/mL; Clinisciences, C311-50ug) and CX3CL1 (100ng/mL; Peprotech, 300-31). At day 41, microglial cells were harvested using Trypsin treatment for 4 min at 37°C. Full recovery of cells was performed by flushing the wells multiple times with DPBS 1X. Cells were collected and centrifuged at 1100rpm for 5min and resuspended in DPBS 1X, following by cell counting (Trypan blue; Life technologies, 15250061).

### Brain organoids production and grafting assays

Human brain organoids (BORGs) were generated by using a modified protocol described by *Rosebrock et al.*^31^ Briefly, iPSCs, at a confluency of 80%, were first singularized by successive exposure to EDTA (0,5mM; Life Technologies AM9262) and Accutase (500U/mL; Sigma-Merck, A6964-100ML). Accutase was inactivated by adding DMEM/F12 (Life technology, 11330057), cells were pipetted slowly to break remaining cell clumps, recovered in DMEM/F12 and centrifuged for 5min at 1000 rpm. Pelleted cells were resuspended in 1mL of embryonic bodies formation medium (Mix 4:1 DMEM/F12 KnockOut™ Serum Replacement KOSR (Life technology, 10828028), 0.5X Glutamax supplement, 1X MEM-NEAA, 1% Penicillin/Streptomycin (Life tech, 15140122), β-mercaptoethanol (Life technology, 31350010). The Basal medium is supplemented, before use, with a cocktail of selective pathway inhibitors: 50µM Rock inhibitor (Tocris, 1254), 4ng/mL BFGF (Peprotech, 100-18B), 10µM SB432542 (Tocris, 1614), 100nM LDN1933189 (StemCell Technologies, 72147), and 3,3µM XAV939 (Tocris, 3748).

9,000 iPS cells, resuspended in EB medium, were grown in 96 well U-bottom Ultra-low attachment plates (150 mL per well; Corning, 7007). At day 7, medium is replaced by 150µL of Neural Induction medium (NIM: DMEM/F12 1X N2, 1X Glutamax Supplement, 1X MEM-NEAA, 1% Penicillin/Streptomycin, supplemented with SB431542, LDN193189, and XAV939). Two days later, EBs were transferred to Organoid Embedding Sheet molds (Stem Cell, 08579) and embedded in 15µL of Growth Factor Reduced Matrigel (Matrigel GFR Corning ref: 354230). Droplets were placed at 37°C for 45min, placed in 6 well Ultra-Low Attachment Plates (Corning, 3471) with 4mL of Expansion medium per well (Expansion Medium: Mix 1:1 DMEM/F12 Neurobasal Medium (Life technology), 1X Glutamax Supplement, 0.5X MEM-NEAA, 0.5X N2, 1X B27 without Vitamin A (Life technologies, 12587010), Insulin (Sigma-Merck 2,5mg/mL, Sigma-Merck, I9278-5ML), 0.05X β-mercaptoethanol). 4 days later, the medium was entirely replaced by 4ml of Maturation Medium (Mix 1:1 DMEM/F12 Neurobasal Medium, 1X Glutamax Supplement, 0.5X MEM-NEAA, 0.5X N2, 1X B27, Insulin (Sigma-Merck Cf = 2,5µg/mL), 0.05X β-mercaptoethanol), shacked continuously (65 rpm; Infors Celltron), by changing 2/3 of the medium every 3-4 days. Culture is performed under constant agitation.

After 1 month of culture, BORGs were grafted with hematopoietic stem cells by re-embedding them in a droplet of 12µL Matrigel Growth factor reduced GFR, this time containing 10,000 HPCs/µL of Matrigel. Matrigel droplets were placed in maturation medium and under agitation. After one or two-months, BORGs were collected for immunofluorescence, RNA isolation, or flow cytometry assays.

### Flow cytometer analysis

HPCs and MGLs were resuspended in labeling buffer (3% FCS/PBS 1X) before staining. HPCs phenotype was analyzed by using the following monoclonal antibodies (mAbs) from BioLegend: mouse anti CD34-APC (clone 581), mouse anti CD43-FITC (clone MEM-59), mouse anti CD45-FITC (clone HI30). MGLs were phenotypically analyzed by using mouse anti CD45-FITC (clone HI30, BioLegend) and rat anti CD11b-APC (clone M1/70, BioLegend) or mouse anti CD11b-BV570 (clone ICRF44, Sony Biotechnology). Before staining, MGLs were previously incubated with human Fc block (BD Bioscience) for 15 min at 4 C, to block or significantly reduce potential non-specific antibody staining. The following isotype control antibodies from BioLegend were also used: APC rat IgG2b, κ (clone RTK4530), APC mouse IgG2b, κ (clone MOPC-21), FITC mouse IgG2b, κ (clone MOPC-21).

Exclusion of dead cells from flow cytometric analysis were performed by adding 7-AAD staining solution (Miltenyi) before acquisition, at the recommended final concentration of 0.525 μg/mL.

Events were acquired on CytoFLEX LX (Beckman Coulter) flow cytometer and the analyses were performed using Kaluza software (Beckman Coulter).

For flow cytometry assays, brain organoid cells were first singularized using the papain dissociation system (Worthington Biochemical,). The presence of MGLs was analysed using the same antibody as the 2D MGL analysis (mouse anti CD45-FITC and rat anti CD11b-APC or mouse anti CD11b-BV570).

### Latex bead uptake assay

Microglia phagocytosis performance was measured by using fluorescent yellow-green latex beads (Sigma-Aldrich; L4655-1ML). 100-500 thousand MGL cells/well were plated in 24 well plates, previously coated with hESC qualified Matrigel, in 500 µL of microglial basal medium (described above) supplemented with 0.1µg/mL/well LPS (Sigma-Aldrich; L4391-1MG). The following day, fluorescent beads were diluted in 5% FCS/PBS 1X and incubated for one hour at 37°C (2 µL of beads were used for each well). Beads were then resuspended in 100 µL DMEM/F12/well, transferred into the wells and the plate was centrifuged at 1100rpm for 3 min before incubation at 37°C for one hour. Cells were collected in both the supernatant and onto the plate by trypsinization. Cells were pelleted at 1100 rpm for 5 min and finally resuspended in 50 µL of labeling buffer (3% FCS/PBS 1X) for acquisition by flow cytometry. As a negative control for beads uptake, we treated some wells 15 min before the addition of the beads, with 15 µg/mL/well cytochalasin D (Sigma-Aldrich, C2618-200UL), an inhibitor of actin polymerization and hence of the uptake phenomenon.

### Inflammation assay

Mature MGL were plated in 6 well plates coated with hESC qualified Matrigel at a concentration of 1×10^6 cells per well in 2 mL of microglial basal medium (described above). Half of the wells were exposed to 0,1µg/mL/well LPS (Sigma-Aldrich) overnight. The following day, cells were collected for RNA extraction. The expression of some pro-inflammatory (IL1beta, TNF-alpha, IL-12), anti-inflammatory (IL10), chemokines (CCL2, CXCL10) was assessed by RT-qPCR. Primers are described in (RNA isolation, Real-time PCR, and RNA sequencing).

### Cell death detection

At days 2 and 5 during MGL differentiation, cell death was assessed by analyzing the cells with the Dead Cell Apoptosis Kit (Invitrogen, V13242). Cells were stained with Alexa® Fluor 488 annexin V and Propodium Iodide (PI), and apoptosis and necrosis were determined by flow cytometric analysis. Basal apoptosis and necrosis were identically determined on untreated cells.

### BORGs Cryosectioning and Immunofluorescence assays

BORGs were fixed in 4% Paraformaldehyde (PFA, Thermo Fischer; 28908) for 30min at room temperature and washed three times with PBS for 5min before being transferred in 1mL of 30% sucrose. The following day, BORGs are placed in OCT (Avantor VWR; 2581596) and frozen at −80°C. OCT blocks were cryo-sectioned at 20µm, and collected on superfrost microscope slides. Sections were permeabilized using 0.1%TRITON X-100 in PBS for 10min at room temperature. Slides were washed 3 times with PBS 1X and then exposed to blocking solution (PBS 1X, 0.1% BSA, 0.1% Triton X-100) for 30min at 4°C. Blocking solution was replaced with a primary antibody solution (PBS 1X, 0.1% BSA, 0.1% Triton X-100) presenting a primary antibody against IBA1 (mouse monoclonal, abcam; ab178846; 1:500) at 4°C overnight. The following day, excess of primary antibody was washed three times with PSB 1X for 5min. Sections were exposed to a secondary antibody (PBS 1X, 0.1% BSA, 0.1% Triton X-100, Donkey anti-Rabbit IgG (H+L) Secondary Antibody, Alexa Fluor 488 (Life technologies, A-21206; 1:1000) during 1h at room temperature in the dark. Slides are washed three times 5min with PBS 1X and mounted with ProLong™ Diamond Antifade containing DAPI (Life Technologies; P36962). Slides were scanned (Microvisionner) using FITC channel for revealing IBA1 positive cells, which were then quantified and classified (ramified and amoeboid-like shapes) by using ImageJ.

### RNA isolation, Real-time PCR, and RNA sequencing

Total RNA was isolated using the RNAeasy Mini kit (Quiagen, 74104). 500ng of the extracted RNA was used for reverse transcription (High-Capacity cDNA RT kit; Life Technologies; 4368814). Transcribed cDNA was diluted ten-fold and used for real-time quantitative PCR (QuantiTect SYBR Green PCR Kit; Quiagen; 204145) performed with the MX3005P thermocycler (Stratagene).

Real-time qPCR was performed for targeting the following transcripts:

GAPDH (Forward: CCCCGGTTTCTATAAATTGAGC; Revere CTTCCCCATGGTGTCTGAG)

OCT4 (Forward: GCTTCAAGAACATGTGTAAGCTG; Reverse AGGGTTTCCGCTTTGCAT)

CD34 (Forward: TCTCCTATCCTAAGTGACATCAAGG; Reverse GAGTTTGCTGGAAATTTCTGTTCT)

CD43 (Forward: GCATGCTGCCAGTGGCTGTGCT; Reverse TGAGCGTGGGCCGACGGCTA)

CD45 (Forward: TGCAAAACTCAACCCTACCC; Reverse TCACATGTTGGCTTAGATGGAG)

CD11b (Forward: AGAACAACATGCCCAGAACC; Reverse GCGGTCCCATATGACAGTCT)

CD11c (Forward: GTGGTGGTGTGATGCTGTTC; Reverse: ATACTGCAGCCTGGAGGAGA)

P2RY12 (Forward: AACTGGGAACAGGACCACTG; Reverse: ACATGAATGCCCAGATGACA)

GRP34 (Forward: ACCAATCATAGCGACCAACC; Reverse: AGGTCTGCAATGGCTACGTT)

TREM2 (Forward: GGTGCCATTTGAGGAACACT; Reverse: TGGCCGACTCACTTCTTCTT)

SLCO2B1 (Forward: GGCATCCAGTTCATGTTCCT; Reverse: CCAAAACTAAGGCGAAGCAG)

KLF2 (Forward: ACCTGTTGTGTGCATTGGAA; Reverse: GGGGAGAGATACCTCCTTGC)

TREM1 (Forward: AAGGAGCCTCACATGCTGTT; Reverse: CACAGTTCTGGGGCTGGTAT)

TNFA (Forward: TCCTTCAGACACCCTCAACC; Reverse: AGGCCCCAGTTTGAATTCTT

IL1B (Forward: GGGCCTCAAGGAAAAGAATC; Reverse: TTCTGCTTGAGAGGTGCTGA)

IL12 (Forward: AAGGAGGCGAGGTTCTAAGC; Reverse: AAGAGCCTCTGCTGCTTTTG)

IL10 (Forward: TGCAAAACCAAACCACAAGA; Reverse: TCTCGGAGATCTCGAAGCAT)

CCL2 (Forward: CCCCAGTCACCTGCTGTTAT; Reverse: TGGAATCCTGAACCCACTTC)

CXCL10 (Forward: AGGAACCTCCAGTCTCAGCA; Reverse: CAAAATTGGCTTGCAGGAAT)

IBA1 (Forward: GCTGAGCTATGAGCCAAACC; Reverse: TCGCCATTTCCATTAAGGTC) SPI1 (Forward: GAAGACCTGGTGCCCTATGA; Reverse: GAAGCTCTCGAACTCGCTGT).

Illumina RNA Sequencing libraries were constructed from 250-500ng of input RNA with the NEBNext Ultra II RNA Library Prep Kit (E7770; E7775). Libraries were sequenced (150-nt pair-end sequencing; Illumina NovaSeq 6000).

### Cut&tag epigenomics assays

Cut&Tag experiments were carried out as previously described^32,33^ with some modifications. Briefly, 250.000 cells were captured with BioMagPlus Concanavalin A (Polysciences, 86057-3) and incubated with the appropriate primary antibody overnight at 4°C (anti-Histone H3(tri methyl K4) antibody (Abcam, ab8580; 1:50); anti-Histone H3(acetyl K27) antibody (Abcam, ab4729; 1:50); anti-Histone H3(tri methyl K9) antibody (Abcam, ab8898; 1:33); anti-Histone H3(tri methyl K27) antibody (Abcam, ab192985; 1:50)). A secondary antibody was incubated at room temperature for 1 hour (Donkey anti-rabbit IgG antibody: Sigma-Aldrich, SAB3700932; 1:100). Fusion protein ProteinA-Tn5 was prepared in-house (plasmid Addgene #124601) by following a previously described protocol ^33^. Purified PA-Tn5 was loaded with the following adapters compatible with Illumina sequencing (hybridized with the complementary Mosaic sequence [PHO]CTGTCTCTTATACACATCT): MOS-universal:

TACACTCTTTCCCTACACGACGCTCTTCCGATCTAGATGTGTATAAGAGACAG MOS-index: GTTCAGACGTGT.GCTCTTCCGATCTAGATGTGTATAAGAGACAG Loaded PA-Tn5 complex (0.16 μM) was incubated for 1 hour at RT, followed by tagmentation step for 2 hours at 37°C. DNA was extracted by phenol/chloroform (Sigma-Aldrich 77617) followed by ethanol precipitation. Finally, the tagmented DNA was amplified using Illumina primers (NEBNext® Multiplex Oligos for Illumina, NEB #7335S) with the following cycling conditions: 98°C for 30s, 20 cycles of 98°C for 10s and 65°C for 75 s; then final extension at 65°C for 5 min and hold at 12°C. DNA libraries were cleaned using 1x volume of SPRIselect beads (Beckman Coulter, B23318) and Illumina sequenced (150-nt pair-end sequencing; Illumina NovaSeq 6000).

### Bioinformatics analyses

Raw fastq files were aligned to GRch38 human genome using Bowtie2 (v2.4.1) under the default parameters. The quality of the aligned reads was verified by Next Generation Sequencing-Control Quality NGS-CQ^34–36^. Aligned reads were associated with coding transcribing regions using FeatureCounts.

Differential gene expression analysis was performed on R using Noiseq 2.38.0 and DESeq2 1.34.0 packages. Read counts per transcript across samples were first quantile normalized, low counts genes were removed from the analysis using the counts per millions method (CPM), and batch effects were corrected with the ARSyN method. Furthermore, differentially expressed transcripts were filtered based on log2fold change and statistic q-value >0.95. Principal component analysis was performed using the “Factominer” package and the 3D representation used the “3D PCA” package.

Heatmaps were obtained using the “pHeatmap” package, and MultiExperimentViewer(MeV). Gene ontology analyses were performed using <u>Enrichr</u>.

Cut&tag histone modification data were aligned to the human genome using Bowtie2 (v2.4.1) under the default parameters. Chromatin state analysis has been performed with ChromHMM (v1.14). Bedgraph files and peak annotations were obtained with MACS2 and visualized within the IGV genome browser (v.2.4.15). Promoter enrichment maps were generated with SeqMiner (v.1.3.4).

### Comparison with Single-cell RNA-seq data issued from Alzheimer’s patients

Gene lists obtained after differential expression analysis were compared with the public single-cell RNA-sequencing data issued from human brain samples described by Brase et al (2023)^18^. Comparison with this public dataset was performed via their graphical online interface “SNARE” (http://web.hararilab.org/SNARE/).

### Data access

All raw datasets generated on this study have been submitted to the NCBI Gene Expression Omnibus (GEO; http://www.ncbi.nlm.nih.gov/geo/) under accession number XXXXX.

## Acknowledgements

We thank all members of the <u>team SysFate</u> for contributing to the discussion of this project, the Genoscope sequencing platform and the "Imaging and Cytometry Core Facility" of Genethon for their technical support. This work was supported by the institutional bodies CEA, CNRS, and Université d’Evry-Val d’Essonne, the Genopole Thematic Incentive Actions funding (ATIGE-2017) as well as the ‘‘Fondation pour la Recherche Medicale’’ (FRM; funding ALZ-201912009904).

## Author contributions

A Aubert: formal analysis, investigation, and methodology. MG Mendoza-Ferri: investigation, and methodology. A Bramoulle: resources and methodology. F Stüder: data curation, software, and formal analysis. BM Colombo: conceptualization, supervision, writing—original draft, review and editing. MA Mendoza-Parra: conceptualization, formal analysis, supervision, funding acquisition, and writing— original draft, reviewing and editing.

## Conflict of Interest and other Ethics Statements

The authors have no competing interests to declare.

