## Supplementary Figures for "PSEN1^M146V^ and PSEN1^A246E^ mutations associated with Alzheimer’s disease impair proper microglia differentiation"

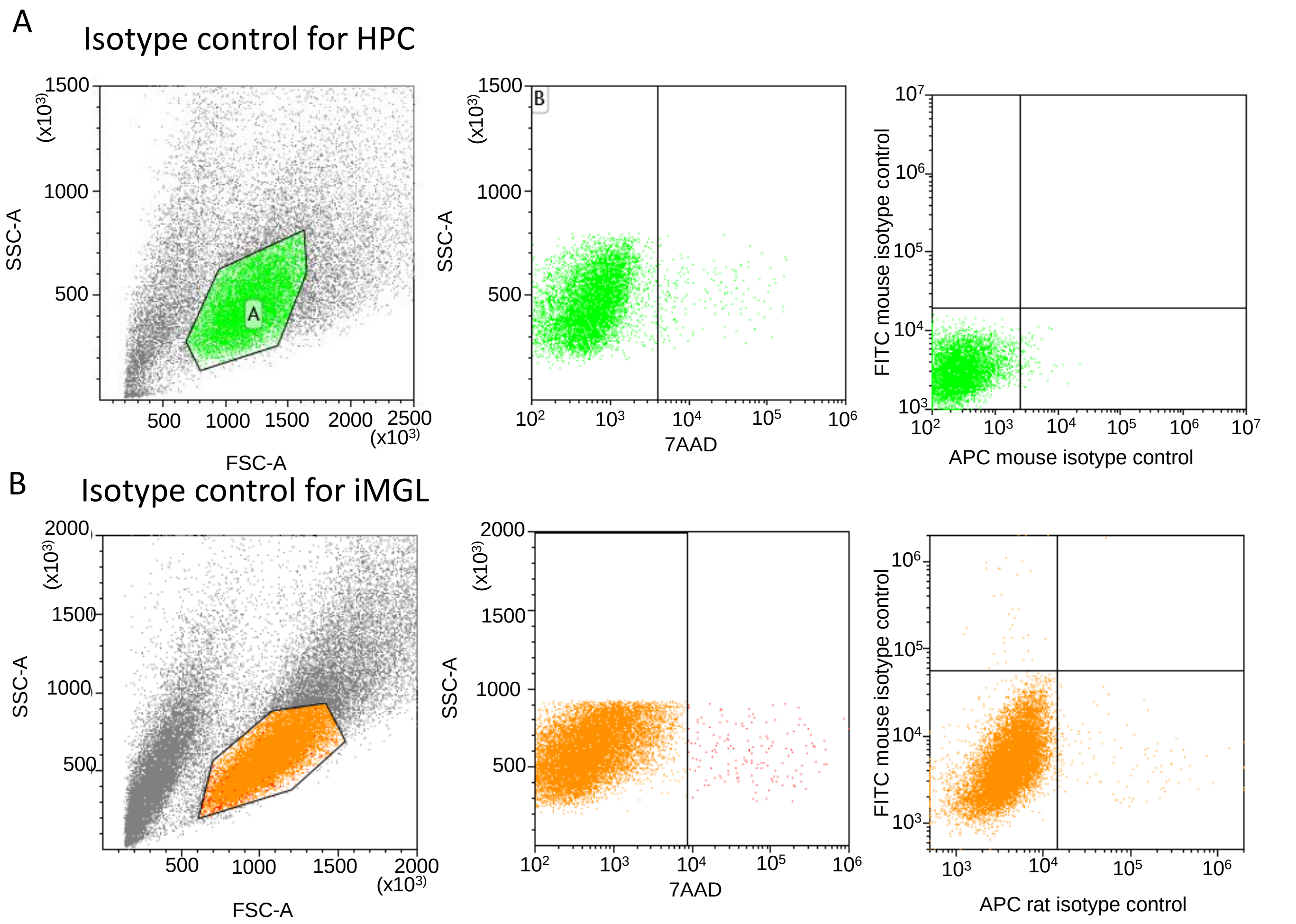


**Extended data Figure 1 related to main Figure 1. Flow cytometry isotype controls.**

**A.** Left panel: Forward Scatter (FSC) versus Side scatter (SSC) plot used for optimal cell gating. Middle panel: Cell viability of the selected population (left panel) was evaluated with 7-AAD staining solution. Right panel: Flow cytometry assay with APC and FITC **i**sotype control antibodies, used as a negative control for the detection of CD34 and CD43 markers for HPCs. **B.** Similar assay than in (A) performed for the detection of the CD11b and CD45 markers, for iMGLs.

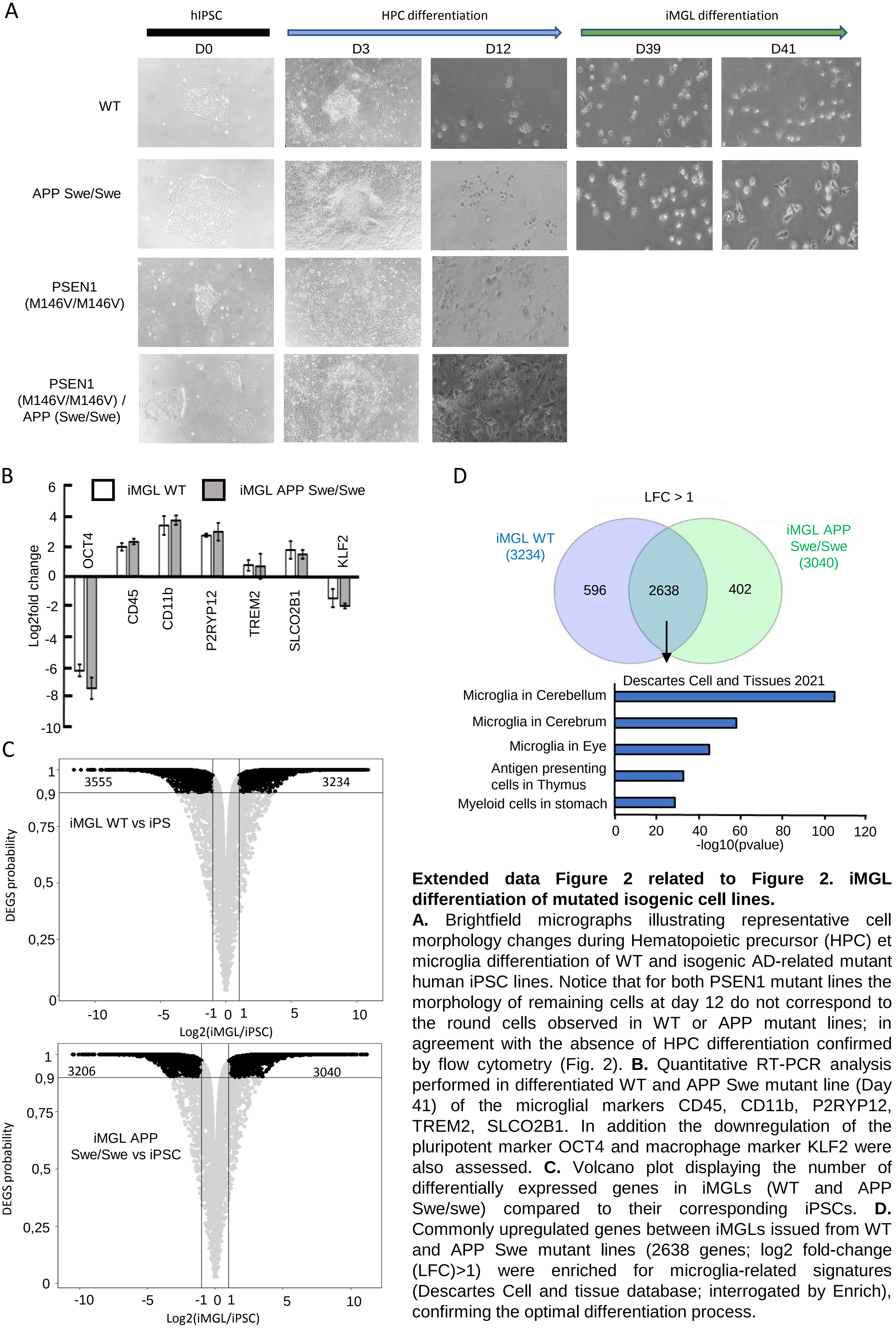


**Extended data Figure 2 related to Figure 2. iMGL differentiation of mutated isogenic cell lines.**

**A.** Brightfield micrographs illustrating representative cell morphology changes during Hematopoietic precursor (HPC) et microglia differentiation of WT and isogenic AD-related mutant human iPSC lines. Notice that for both PSEN1 mutant lines the morphology of remaining cells at day 12 do not correspond to the round cells observed in WT or APP mutant lines; in agreement with the absence of HPC differentiation confirmed by flow cytometry (Fig. 2). **B.** Quantitative RT-PCR analysis performed in differentiated WT and APP Swe mutant line (Day 41) of the microglial markers CD45, CD11b, P2RYP12, TREM2, SLCO2B1. In addition the downregulation of the pluripotent marker OCT4 and macrophage marker KLF2 were also assessed. **C.** Volcano plot displaying the number of differentially expressed genes in iMGLs (WT and APP Swe/swe) compared to their corresponding iPSCs. **D.** Commonly upregulated genes between iMGLs issued from WT and APP Swe mutant lines (2638 genes; log2 fold-change (LFC)>1) were enriched for microglia-related signatures (Descartes Cell and tissue database; interrogated by Enrich), confirming the optimal differentiation process.

A

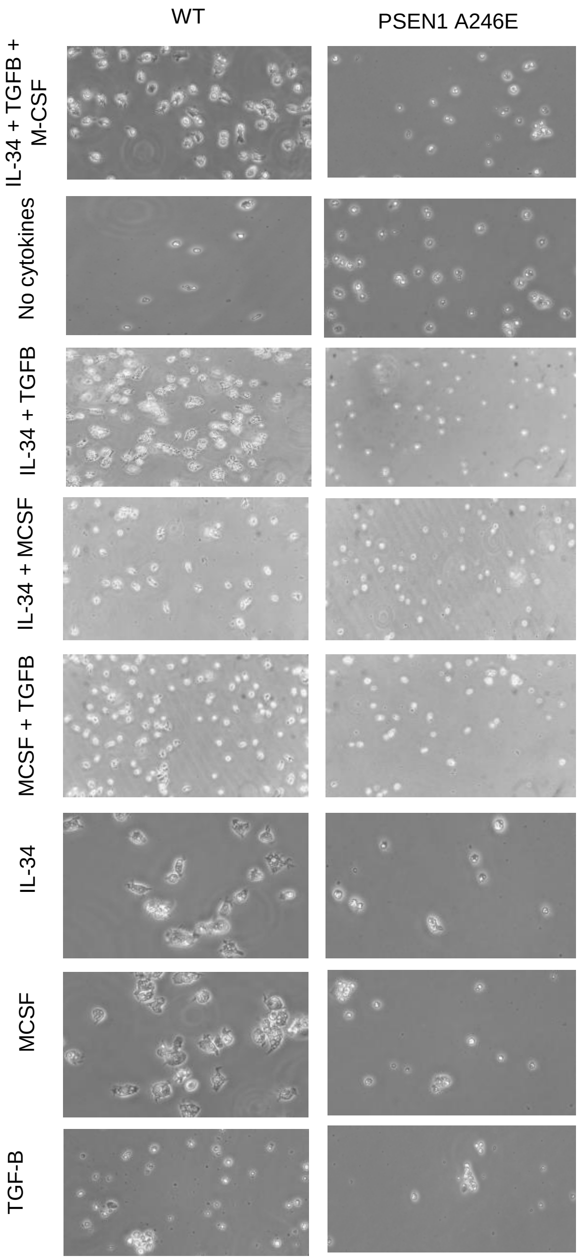


B

Microglia diff. induction

| Cytokine drivers | days | WT<br>Annexin V+/PI+ cells | PSEN1 (A246E/WT)<br>Annexin V+/PI+ cells |
|---|---|---|---|
| **IL-34 + TGF-B + M-CSF** | **D2** | **20,96%** | **84,70%** |
| | **D5** | **9,25%** | **98,70%** |
| **Absence of cytokine** | **D2** | **58,70%** | **84,46%** |
| | **D5** | **87,13%** | **98,84%** |
| IL-34 + TGF-B | D2 | 55,51% | 87,87% |
| | D5 | 49,01% | 99,08% |
| IL-34 + M-CSF | D2 | 54,56% | 87,61% |
| | D5 | 44,27% | 99,16% |
| TGF-B + M-CSF | D2 | 53,86% | 86,80% |
| | D5 | 13,76% | 98,93% |
| IL-34 | D2 | 27,08% | 86,60% |
| | D5 | 72,35% | 98,95% |
| TGF-B | D2 | 65,37% | 87,71% |
| | D5 | 93,07% | 99,06% |
| M-CSF | D2 | 24,80% | 84,15% |
| | D5 | 81,30% | 99,09% |

**Extended data Figure 3 related to Figure 2. Cell viability assay in presence of different cytokine combinations required for microglia differentiation induction.**

**A.** Brightfield micrographs displaying representative HPC morphology issued from either WT or PSEN1 A246E mutant line, after 5 days of treatment with the indicated cytokine combination. **B.** Percentage of Apo-Necrotic (Annexin V+ / Propidium Idodide PI+) cells after 2 and 5 days in the presence of the indicated cytokine combination.

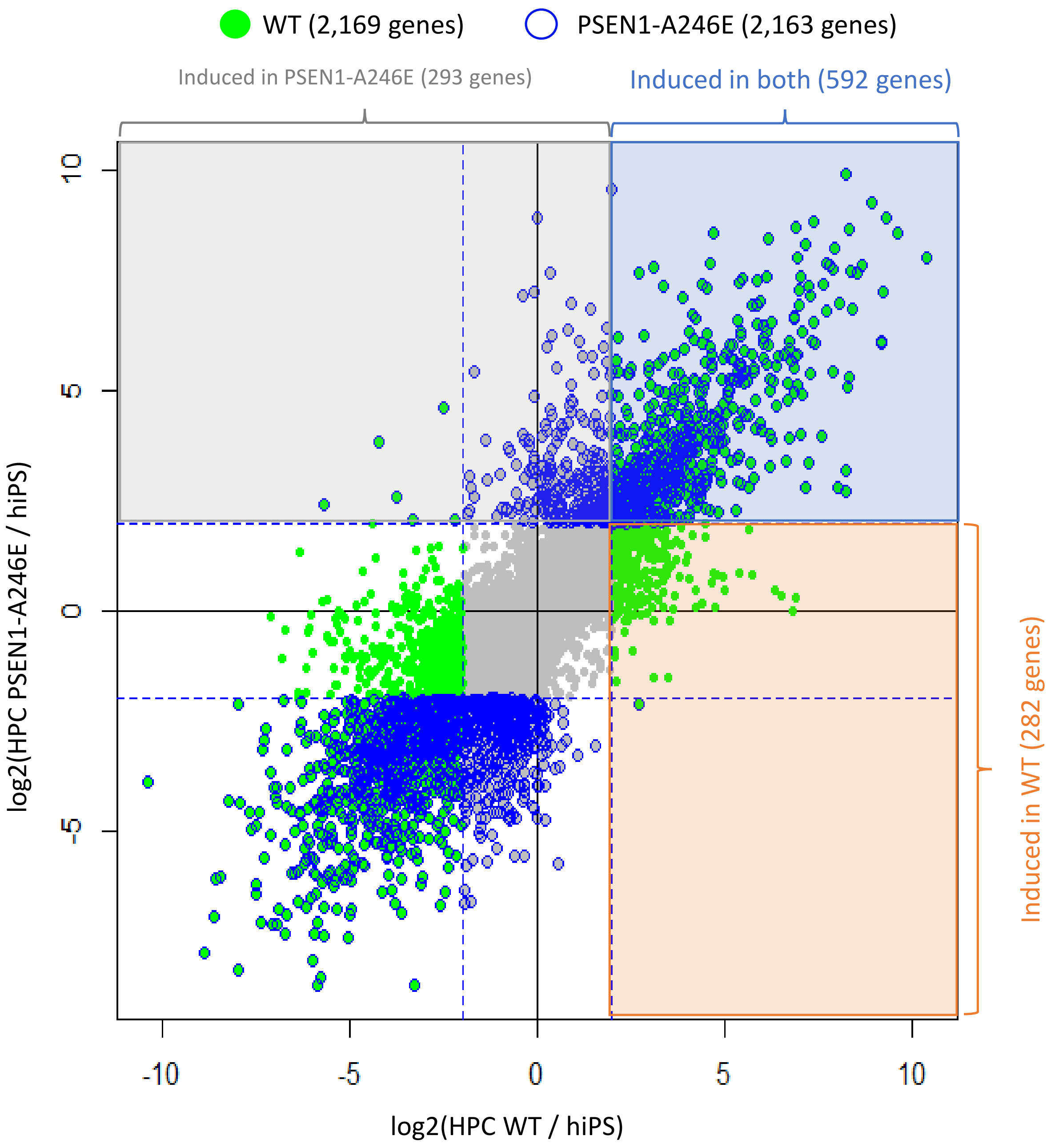


**Extended data Figure 4 related to Figure 3. Changes in gene expression between HPCs issued from WT and PSEN1 A246E mutant cells.**

Scatter-plot displaying a comparison between the differential expression levels (relative to their corresponding hiPSCs) assessed in WT HPCs (X axis) and HPCs harboring the PSEN1-A246E mutation (Y axis). Gene were considered differently expressed with a log2fold change > |3|. Up-regulated genes were classified in three groups: (i) induced in both lines (592 genes), (ii) Induced only in WT HPCs (282 genes), (iii) Induced only in the PSEN1-A246E mutant HPC (293 genes).

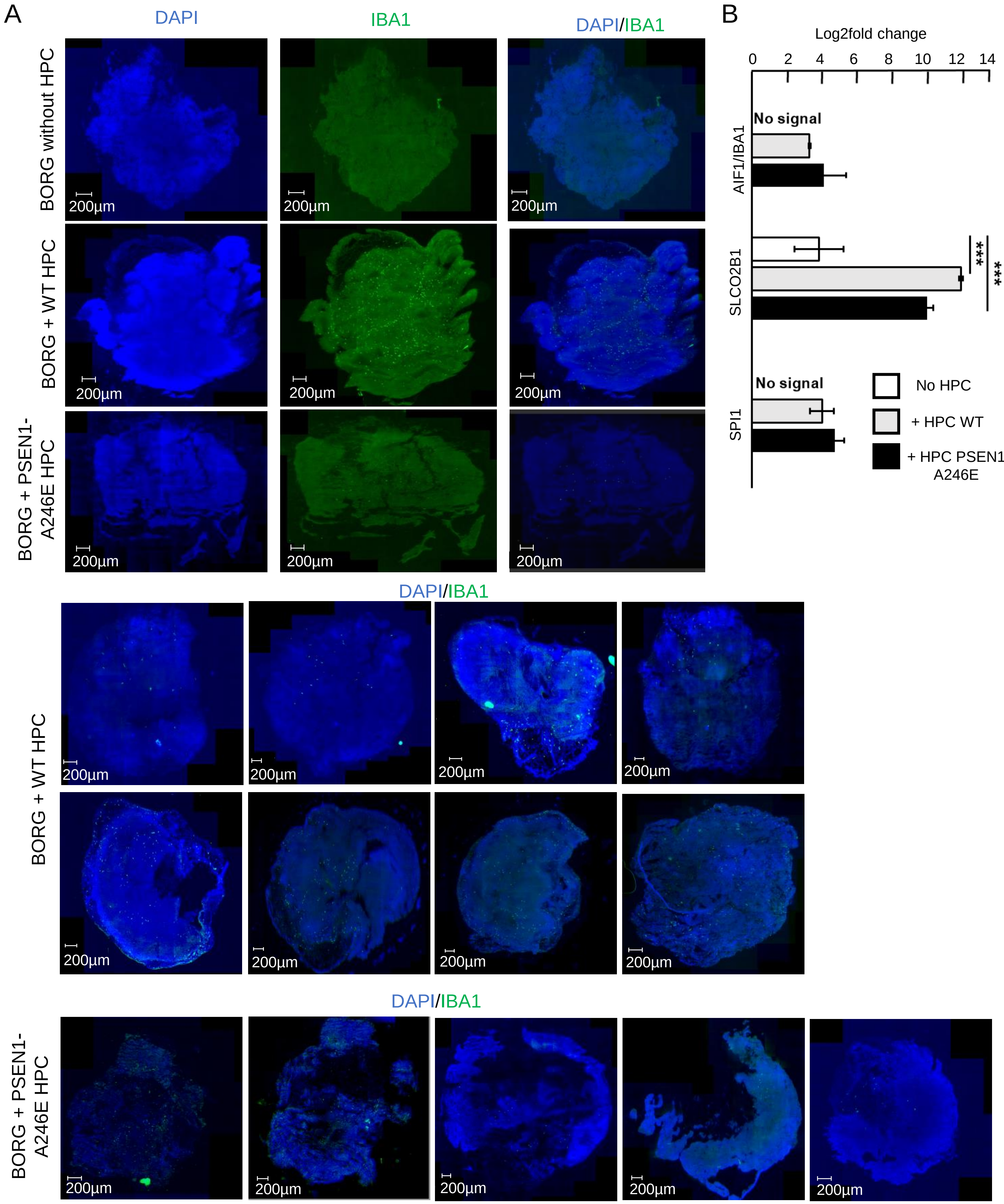


**Extended data Figure 5 related to Figure 4 : Evaluation of the presence of microglial cells in brain organoids grafted with HPCs.**

**A.** Manually scanned micrographs revealing nuclei (DAPI) and the microglia marker IBA1 in either ungrafted BORGs or graffted BORG with HPC issued from WT or mutant PSEN1-A246E mutant hiPSCs. Immunolabelling assay has been performed two months after grafting. **B.** Quantitative RT-PCR analysis targeting the microglial markers IBA1, SCLO2B1 or SPI1.

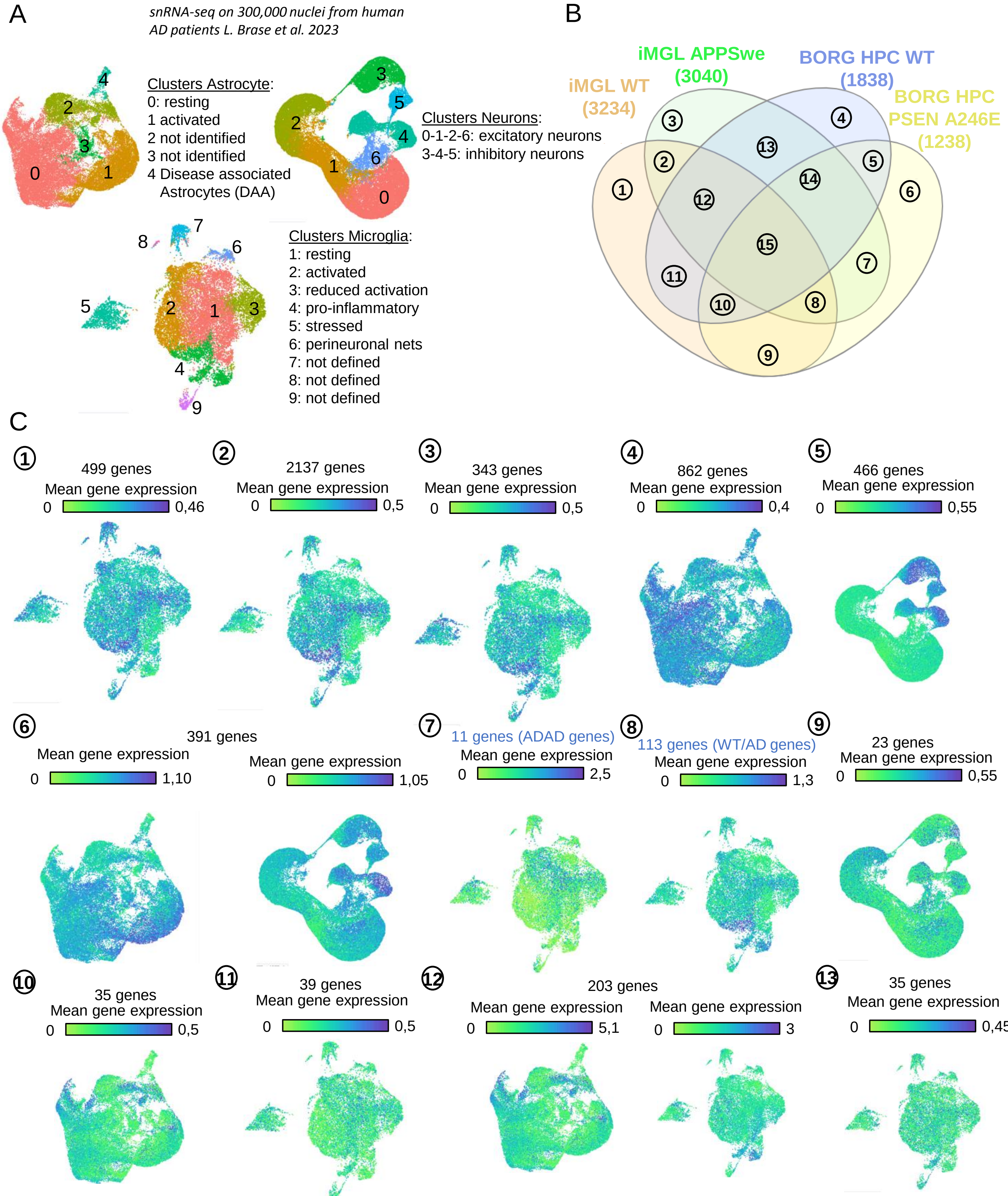

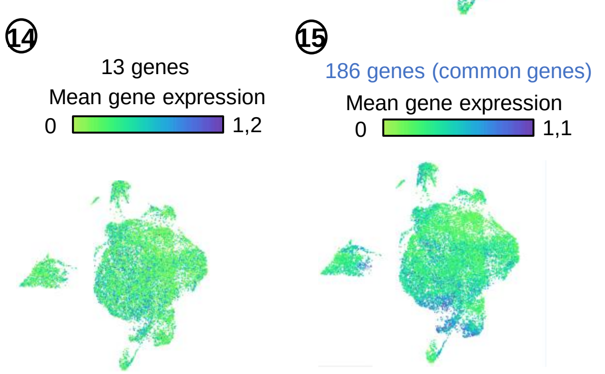


**Extended data Figure 6 related to Figure 5. Upregulated genes retrieved in in-vitro generated microglia from AD-related mutant lines correlate with microglial signatures observed in human brain samples issued from AD patients. A.** UMAP Clustering of single cell RNA-seq datasets performed by Brase et al 2023. **B.** Venn-diagram illustrating the number of up-regulated genes in four microglial differentiation conditions described in this study. **C.** Mean gene expression signatures, within the UMAP clusters displayed in (A), associated to each of the up-regulated intersection groups observed in (B).
